# The arginine/ornithine binding protein ArgT plays an essential role in *Brucella* to prevent intracellular killing and contribute to chronic persistence in the host

**DOI:** 10.1101/2023.07.11.548583

**Authors:** Sushree Rekha Mallik, Kiranmai Joshi, Girish K. Radhakrishnan

## Abstract

*Brucella* species are facultative intracellular bacterial pathogens that cause the contagious zoonotic disease, brucellosis. *Brucella* spp. infect a wide range of animals, including livestock, wild animals, and marine mammals. Brucellosis remains endemic to various parts of the world, affecting the economic growth of many countries because of its impact on public health and livestock productivity. There are no human vaccines for brucellosis, and controlling the disease in susceptible animals is crucial for limiting human infections. Although the available live-attenuated vaccines have protective efficacy in animals, they have many disadvantages, including infectivity in humans. Compared with other invasive bacterial pathogens, minimal information is available on the virulence factors of *Brucella* that enable them to survive in the host. Here, we performed transposon-based random mutagenesis of *B. neotomae* and identified the arginine/ornithine binding protein, ArgT, as the crucial virulence determinant of *Brucella*. Deleting *ArgT* from *B. melitensis* resulted in its attenuation in macrophages, which was restored upon complementation with an *ArgT* expression plasmid. We observed that macrophages infected with Δ*ArgT*-*Brucella* produced elevated levels of NO due to the inability of Δ*ArgT Brucella* to deplete the host intracellular arginine through its importer. Furthermore, defective survival of Δ*ArgT B. melitensis* was observed in the infected mice, which correlated with enhanced NO production in the mice. Our studies revealed that *ArgT* in *Brucella* plays a vital role in preventing intracellular killing and contributes to the chronic persistence of *Brucella* in the host. This study highlights the essential role of arginine in clearing intracellular infections and the subversion of this host defense mechanism by intracellular pathogens for their chronic persistence.

## INTRODUCTION

Brucellosis is a zoonotic disease worldwide that accounts for more than 500,000 new human infections annually (1). Brucellosis is endemic to many parts of the developing world, posing a serious threat to the human healthcare system and constraining the economic growth of many countries (2). Despite various control measures, brucellosis remains a major veterinary and public health problem in Asia, Africa, Latin America, the Middle East, and Europe’s Mediterranean and Southeast regions (3). Brucellosis is caused by facultative intracellular gram-negative bacteria belonging to the genus *Brucella. Brucella* infection in domestic animals results in abortion, sterility, reduced milk yield, and weakened offspring. The symptoms of human brucellosis include undulating fever, chills, headaches, and joint pain (4). Chronic brucellosis can lead to various complications, such as arthritis, epididymal orchitis, endocarditis, splenic abscess, abortion, and neurobrucellosis (5). Twelve species of *Brucella* have been identified with distinct host specificity, and four *Brucella* species, *B. melitensis, B. abortus, B. suis*, and *B. canis*, have been reported to cause human infections (6). Humans can contract the disease by ingesting contaminated dairy products, directly contacting infected animals, or inhaling *Brucella*-containing aerosols. Currently, there are no licensed vaccines for human brucellosis. Antibiotic treatment for human brucellosis is less efficient because of the prolonged duration of treatment with a combination of antibiotics, frequent relapses, and therapeutic failures (7). Available animal vaccines such as *B. abortus* S19, RB51, and *B. melitensis* Rev1 are effective. However, they have many disadvantages, including infectivity in humans, induction of abortion in pregnant animals, and low protection efficacy (8). Therefore, it is essential to develop improved therapeutic and preventive strategies for animal and human brucellosis.

*Brucella* can survive and replicate in both professional and non-professional phagocytic cells. *Brucella* exploits various host cellular processes to invade and replicate in phagocytic cells, thus making it a successful intracellular pathogen. *Brucella* enters host cells through lipid rafts, resists intracellular killing, and establishes a permissive replication niche (9). *Brucella* modulates endocytic trafficking pathways to avoid the fusion of *Brucella*-containing vacuoles with lysosomes and eventually fuses with the endoplasmic reticulum to form a replicative niche (10). *Brucella* secretes effector proteins through its Type IV Secretory System to modulate host responses for replication and chronic persistence (11). However, there is minimal information on *Brucella* effector proteins and their host targets. Identifying the mechanisms by which *Brucella* subverts host cellular processes can lead to the development of improved vaccines and therapeutics for brucellosis.

Transposable genetic elements have been widely employed as molecular biology tools to introduce insertions into genes to generate mutations (12). Here, we performed transposon mutagenesis of *B. neotomae* which mimics the infection dynamics of its zoonotic counterpart, *B. melitensis*. Mutagenesis screening identified the gene coding for the arginine/ornithine binding protein ArgT as the crucial determinant of *Brucella* survival in macrophages. Arginine plays a vital role in the host defense against infectious microbes as it can serve as a source of nitric oxide (NO). Activated macrophages undergo two major types of polarization, *viz*. M1 and M2 (13). M1 macrophages exhibit microbicidal properties and produce pro-inflammatory cytokines, whereas M2 macrophages exhibit anti-inflammatory and tissue-repair properties (13). M1 macrophages express nitric oxide synthase (NOS), which metabolizes arginine to NO and citrulline. NO plays a vital role in the microbicidal properties of macrophages, and citrulline is recycled to more NO production via the citrulline-NO cycle (14). The intracellular availability of arginine determines the NO production. In contrast, M2 macrophages express arginase, which hydrolyzes arginine into ornithine and urea, which can contribute to tissue repair and cell proliferation by producing polyamines and proline from ornithine (15). Arginase can limit NO production in M2 macrophages by hydrolyzing arginine (14).

Pathogenic microorganisms have evolved strategies to deplete the host arginine pool and attenuate NO production, which facilitates their intracellular survival. They actively transport arginine from the intracellular milieu to their cells through polar amino acid importers, and ArgT plays a major role in arginine binding (16). The imported arginine can be hydrolyzed into ornithine and urea by pathogen-encoded arginase (17). This depletes the intracellular availability of arginine, leading to attenuation of NO production. We found that Δ*ArgT*-*Brucella* presented an attenuated phenotype in macrophages and mice. We observed enhanced production of NO in macrophages and in mice infected with Δ*ArgT B. melitensis*. Our study revealed that *ArgT* plays an essential role in the intracellular survival of *Brucella* by attenuating the production of NO in infected cells and animals. This information may help develop novel vaccines and therapeutics targeting the arginine/ornithine transporter in *Brucella* and other invasive bacterial pathogens.

## RESULTS

### *ArgT* is essential for the intracellular survival of *Brucella* in macrophages

While performing random mutagenesis of *B. neotoma*e using *Himar1* transposon (Tn), we identified a mutant that exhibited defective survival in macrophages. We infected RAW264.7 with Tn mutant *B. neotomae* and analyzed the intracellular load of bacteria at various times post-infection. We observed defective survival of *B. neotomae* mutant at later time points in macrophages compared to the wild type (WT) (Fig. 1A). To further confirm the experimental data, we introduced a Green Fluorescence Protein (GFP) expression plasmid into WT or Tn mutant *B. neotomae*, followed by macrophage infection studies. RAW264.7 cells were infected with GFP-expressing *Brucella*, followed by analysis of the infected cells 24 h post-infection using a fluorescent microscope. In agreement with the CFU analysis, a diminished number of mutant *B. neotomae* was observed in macrophages compared to the WT (Fig. 1B). These studies suggest that Tn mutant *B. neotomae* can invade and replicate in macrophages. However, they exhibit defective intracellular survival at later stages of infection.

**Figure 1.**
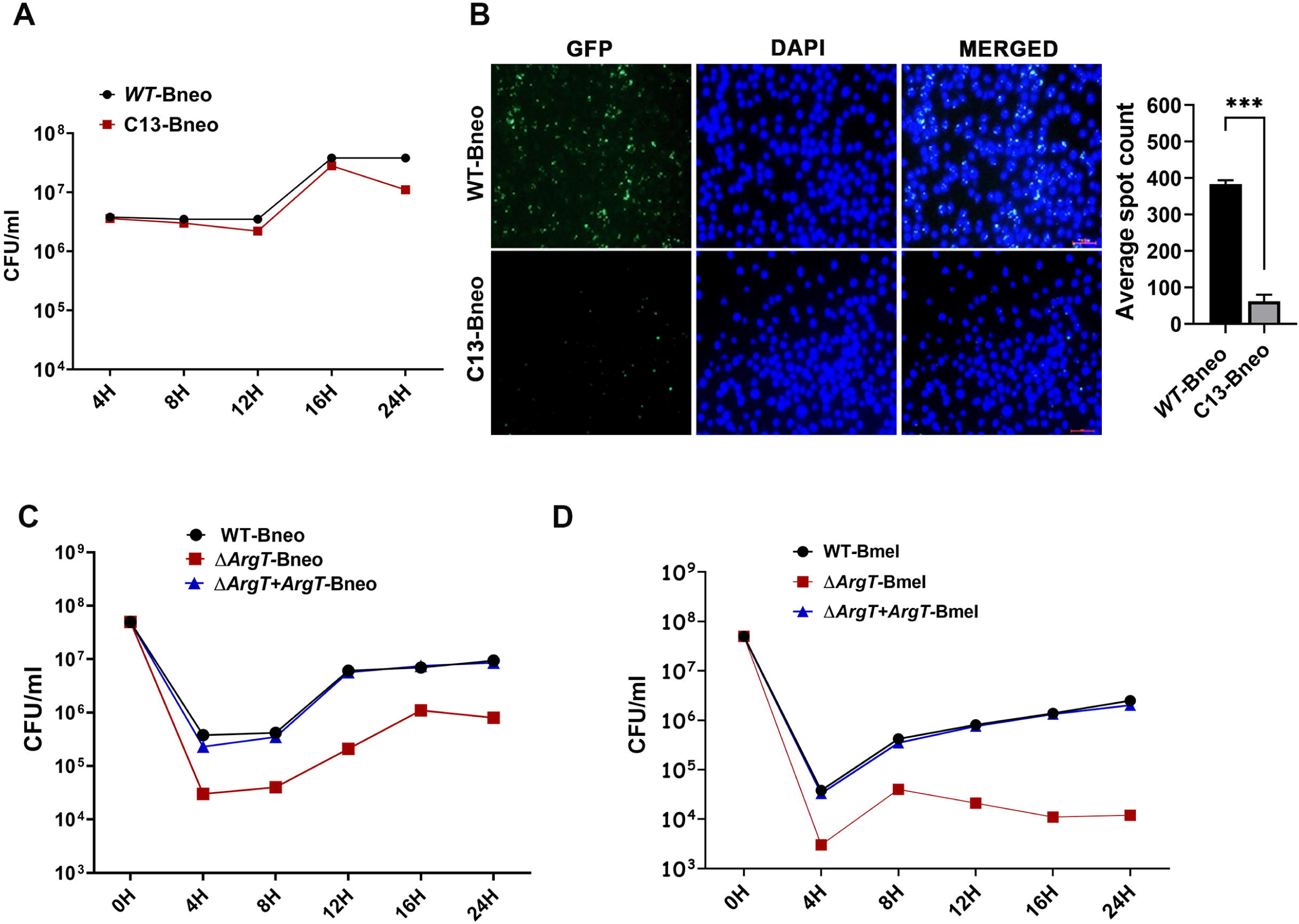
Δ*ArgT Brucella* exhibits defective intracellular survival in the macrophages. **(A)** RAW264.7 cells were infected with wild-type (WT-Bneo) or Transposon-insertional mutant of *B. neotomae* (C13-Bneo), followed by quantification of the intracellular load of bacteria by enumeration of CFU at indicated times post-infection. **(B)** Fluorescence microscopy images showing the intracellular load of WT-Bneo or C13-Bneo expressing GFP. RAW264.7 cells were infected with GFP-expressing *Brucella*, followed by analysing the infected cells using a fluorescent microscope at 24 hours post-infection. The cell nuclei were stained with DAPI (blue). Scale bar, 50 µm. The images are representative of three independent experiments where 12 fields were captured and analysed for each experiment. The right panel indicates the quantification of the number of intracellular bacteria using ImageJ software. **(C)** Macrophage infection studies using WT (WT-Bneo) or Δ*ArgT*-*B. neotomae* (Δ*ArgT*-Bneo) *or* Δ*ArgT*-*B. neotomae* complemented with *ArgT* expression plasmid (Δ*ArgT+ArgT*-Bneo). RAW264.7 cells were infected with indicated strains of *B. neotomae*, followed by the enumeration of CFUs at various times post-infection. (**D**) The intracellular load of WT (WT-BmeI) or Δ*ArgT B. melitensis* (Δ*ArgT*-BmeI) or Δ*ArgT+ArgT B. melitensis* (Δ*ArgT+ArgT*-BmeI) in RAW264.7 cells at indicated times post-infection. The data are presented as the mean ± SEM from at least three independent experiments (***, PL<L0.001).

Next, we sought to identify the gene disrupted by transposon insertion in the mutant *B. neotomae*. To examine this, the genomic DNA of the Tn mutant *B. neotomae* was isolated, followed by partial digestion with *Hha*1 restriction enzyme and self-ligation to obtain circular DNA fragments (Suppl. Fig. 1B). Subsequently, the ligated product was used as a template for inverse PCR using primers from the kanamycin-resistance (KanR) cassette to amplify the junction at which the transposon was integrated (Suppl. Fig. 1C). The PCR amplicon was cloned and sequenced to identify target gene sequences (Suppl. Fig. 1D). Sequence analysis indicated that the gene disrupted by *Himar1* transposon was homologous to BMEI1022 of *B. melitensis*, which has been reported as an arginine/ornithine-binding periplasmic protein precursor (*ArgT*) (Suppl. Fig. 1E). The *ArgT* belongs to the ATP-Binding Cassette (ABC) transporter complex, where it binds to arginine to transport it through the importer (18).

To examine the role of *ArgT* in *Brucella* virulence of *Brucella*, we generated Δ*ArgT*-*B. neotomae* and *B. melitensis*, followed by macrophage infection studies. A homologous recombination-based gene replacement technique was used to replace *ArgT* with a KanR cassette. Subsequently, Δ*ArgT*-*B. neotomae* and *B. melitensis* were confirmed by PCR analysis (Suppl. Fig. 2). We also generated Δ*ArgT+ ArgT*-*B. neotomae* and *B. melitensis* strains harboring an *ArgT* expression plasmid to complement the *ArgT* deficiency. To examine the infection dynamics of *Brucella* mutants, RAW264.7 were infected with WT, Δ*ArgT* or Δ*ArgT+ ArgT*-*Brucella*, followed by analysis of their intracellular replication at various times post-infection. In agreement with previous observations, both Δ*ArgT-B. neotomae* and *B. melitensis* exhibited defective intracellular survival at later time points. However, this deficiency was restored in Δ*ArgT+ArgT-Brucella*, confirming the specific role of *ArgT* in intracellular survival of *Brucella* (Fig. 1C and 1D).

### Production of NO is enhanced in the macrophages infected with **Δ***ArgT*-*Brucell*a

Intracellular arginine acts as a source of NO production to kill microbial pathogens that are phagocytosed or invaded. However, microbial pathogens deplete the intracellular availability of arginine through active cellular uptake, which can minimize NO production. To examine whether Δ*ArgT*-*Brucella* was defective in utilizing arginine, WT, Δ*ArgT* or Δ*ArgT + ArgT-B. neotomae* were inoculated into arginine-deficient or arginine-supplemented media, followed by measurement of OD at 600 nm at various times post-inoculation. We observed poor growth in WT, Δ*ArgT* and Δ*ArgT+ArgT*-*B. neotomae* in arginine-deficient media (Fig. 2A). However, the growth defect was restored in WT and Δ*ArgT+ArgT*-*B. neotomae* when the arginine-deficient media were supplemented with L-arginine. In contrast, Δ*ArgT B. neotomae* exhibited poor growth in arginine-supplemented media, indicating its inability to uptake arginine into the bacteria due to the deficiency of *ArgT*.

**Figure 2.**
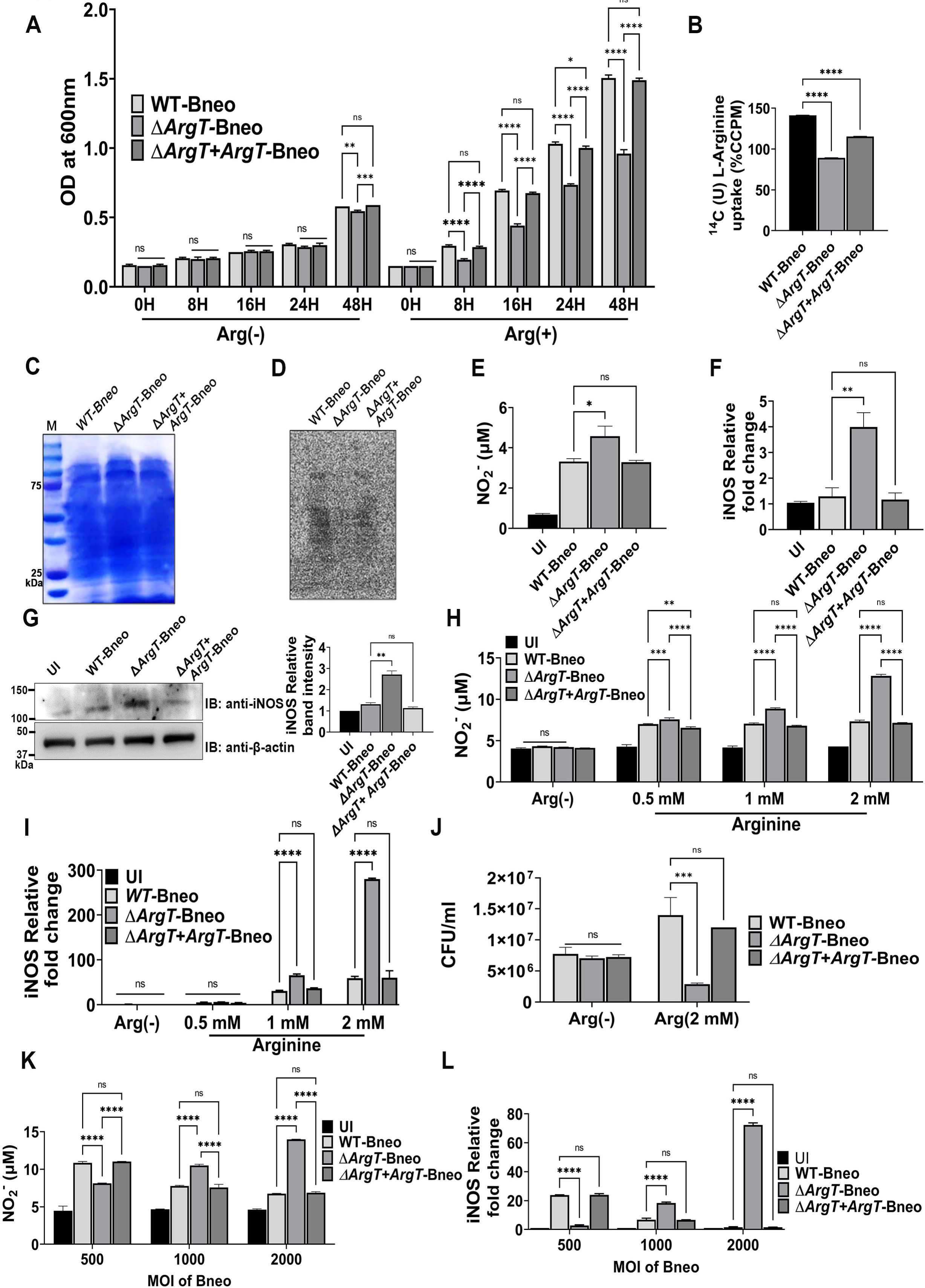
(A) Δ*ArgT*-*B. neotomae* exhibits a defective arginine uptake. Growth dynamics of WT (WT-Bneo) or Δ*ArgT*-*B. neotomae* (Δ*ArgT*-Bneo) or Δ*ArgT+ArgT*-*B. neotomae* (Δ*ArgT*+*ArgT*-Bneo) in the media with or without arginine. The indicated *B. neotomae* strains were inoculated into the arginine (-) or arginine (-) media supplemented with L-arginine. The OD was analyzed at indicated times post-inoculation**. (B) The uptake of C^14^-labeled L-arginine by wild-type or mutant strains of *B*. neotomae**. The uptake assay was performed with WT-Bneo or Δ*ArgT*-Bneo or Δ*ArgT+ArgT*-Bneo in the presence of ^14^C(U) L-arginine. Subsequently, the bacteria were lysed, and the imported ^14^C(U) L-arginine was quantified using the liquid scintillation counter **(C, D) Analysis of incorporated ^14^C(U) L-arginine in the *Brucella* proteins.** After the ^14^C(U) L-arginine uptake assay, WT-Bneo or Δ*ArgT*-Bneo or Δ*ArgT+ArgT*-Bneo were cultured for 24 hours, followed by lysing the cells and resolving equal amount of total protein by SDS-PAGE **(C).** The gel was then dried and analysed using a phosphor-imager **(D)**. **(E)** Δ***ArgT-*B.neo-infected macrophages produce elevated levels of NO.** RAW264.7 cells were infected with WT-Bneo or Δ*ArgT*-Bneo or Δ*ArgT+ArgT*-Bneo, followed by quantification of NO levels at 24 hours post-infection**. (F&G) Enhanced level of iNOS expression in the macrophages infected with** Δ***ArgT*-Bneo.** RAW264.7 cells were infected with WT-Bneo or Δ*ArgT*-Bneo or Δ*ArgT+ArgT*-Bneo, followed by quantification of iNOS mRNA levels by qPCR **(F)** and iNOS protein levels by immunoblotting **(G)** at 24 hours post-infection. The membranes were probed with anti-iNOS antibody to detect the endogenous levels of iNOS. Actin served as the loading control. The right panel indicates the densitometry of iNOS bands normalized with actin. The immunoblot is representative of three different experiments. **(H -J) The Addition of L-arginine enhanced host-induced NO and iNOS levels in macrophages.** RAW264.7 cells were infected with WT-Bneo or Δ*ArgT*-Bneo or Δ*ArgT+ArgT*-Bneo in DMEM medium (-) arginine or arginine (-) media supplemented with indicated concentration of L-arginine, followed by quantification of NO levels in the culture supernatants **(H)**, iNOS expression levels in the cells **(I)**, and determination of the intracellular load of *B. neotomae* strains at 24 hours post-infection **(J)**. **(K & L) The levels of NO and iNOS in the macrophages infected with various MOIs of *Brucella*.** RAW264.7 cells were infected with an increasing MOI of WT-Bneo or Δ*ArgT*-Bneo or Δ*ArgT+ArgT*-Bneo, followed by quantification of NO levels in the culture supernatants **(K)** and iNOS expression levels in the cells **(L)** at 24 hours post-infection. The data are presented as the mean ± SEM from at least three independent experiments (*, P <L0.05; **, PL<L0.01; ***, PL<L0.001; ****, PL<L0.0001).

To validate these data further, we performed an arginine uptake assay using radiolabeled ^14^C(U) L-arginine. Wild-type or Δ*ArgT or* Δ*ArgT*+*ArgT*-*B. neotomae* were cultured in an arginine-deficient medium supplemented with ^14^C(U) L-arginine, followed by measurement of radiolabeled arginine in *Brucella* lysates by liquid scintillation counting. We observed diminished uptake of ^14^C(U) L-arginine by Δ*ArgT*-*B. neotomae* compared to the WT or *B.* Δ*ArgT*+*ArgT*-*B. neotomae* (Fig. 2B). Furthermore, the ^14^C(U)-L-arginine uptake assay was performed with WT, Δ*ArgT or* Δ*ArgT*+*ArgT*-*B. neotomae*, followed by culturing the *Brucella* strains for 24 h to incorporate imported radiolabeled ^14^C(U) L-arginine into their proteins. Subsequently, the bacterial pellets were lysed and subjected to SDS-PAGE to analyze the incorporation of radiolabeled ^14^C(U) -arginine using a phosphor-imager. We observed a decreased level of ^14^C(U) L-arginine incorporation in Δ*ArgT B. neotomae* compared to that in WT or Δ*ArgT*+*ArgT*-*B. neotomae* (Fig.2C and D). The experimental data suggested that *Brucella ArgT* plays an essential role in arginine import.

Next, we examined the levels of NO in macrophages infected with WT or mutant *B. neotomae*. RAW264.7 cells were infected with WT, Δ*ArgT*, or Δ*ArgT + ArgT*-*B. neotomae*. Twenty-four hours post-infection, we analyzed NO production by estimating the nitrite released into the culture supernatant. We observed elevated NO levels in macrophages infected with Δ*ArgT*-*B. neotomae* compared to WT or Δ*ArgT +ArgT*-*B. neotomae* (Fig. 2E). To further confirm the data, we analyzed the expression level of iNOS in infected macrophages, which produces NO from arginine. RAW264.7 cells were infected with WT or mutant *B. neotomae* for 24 h, followed by analysis of iNOS levels by qPCR and immunoblotting. Enhanced iNOS expression was observed in macrophages infected with Δ*ArgT*-*B. neotomae* compared with the WT or Δ*ArgT+ArgT* strains (Fig. 2F and G). As indicated previously, we observed a diminished load of intracellular Δ*ArgT*-*B. neotomae* 24 h post-infection, which correlates with the enhanced production of NO (Fig. 1C).

To further confirm the experimental data, we analyzed the production of NO in *B. neotomae*-infected macrophages treated with increasing concentrations of L-arginine. RAW264.7 macrophages were treated with various concentrations of L-arginine, followed by infection with WT, Δ*ArgT* or Δ*ArgT+ ArgT*-*B. neotomae*. iNOS levels and NO production were significantly enhanced in the Δ*ArgT*-*B. neotomae*-infected cells with increasing concentrations of L-arginine. In contrast, WT or Δ*ArgT+ArgT*-*B. neotomae* showed similar levels of NO production and iNOS expression in infected cells (Fig. 2H and I). Intracellular load of Δ*ArgT*-*B. neotomae* declined with increasing concentrations of L-arginine, indicating a direct correlation between the production of NO and its effect on the intracellular survival of *Brucella* (Fig. 2J). Next, we infected RAW264.7, cells with an increasing MOI (Multiplicity Of Infection) of WT or mutant *B. neotomae*, followed by analysis of NO and iNOS levels. Increasing the MOI of Δ*ArgT*-*B. neotomae* enhanced NO production and iNOS expression, whereas WT or Δ*ArgT+ ArgT*-*B. neotomae* efficiently suppressed the production of NO and expression of iNOS in infected cells (Fig. 2K and L).

Given that *Brucella* imports arginine from infected cells, we examined whether *ArgT* and arginase encoded by *rocF* are upregulated in *Brucella* in infected macrophages. To investigate this, we examined the expression levels of *Brucella ArgT* and *rocF* at various time points post-infection. RAW264.7 cells were infected with *B. neotomae*, followed by analysis *ArgT* and *rocF* expression using qPCR. We observed elevated levels of *ArgT* and *rocF* expression at later stages of infection in *B. neotomae* in infected macrophages (Fig. 3A and B). To further confirm these data, we analyzed the expression levels of *ArgT* and *rocF* at various concentrations of L-arginine. RAW264.7 macrophages were treated with increasing concentrations of L-arginine, followed by infection with *B. neotomae* and analysis of the expression of *ArgT* and *rocF* by qPCR. We observed enhanced *ArgT* and *rocF* expression with increasing L-arginine concentrations (Fig. 3C and D). Collectively, our experimental data suggest that *ArgT* and *rocF* are induced in *Brucella* to deplete host arginine and utilize the imported arginine to convert it into other metabolites.

**Figure 3.**
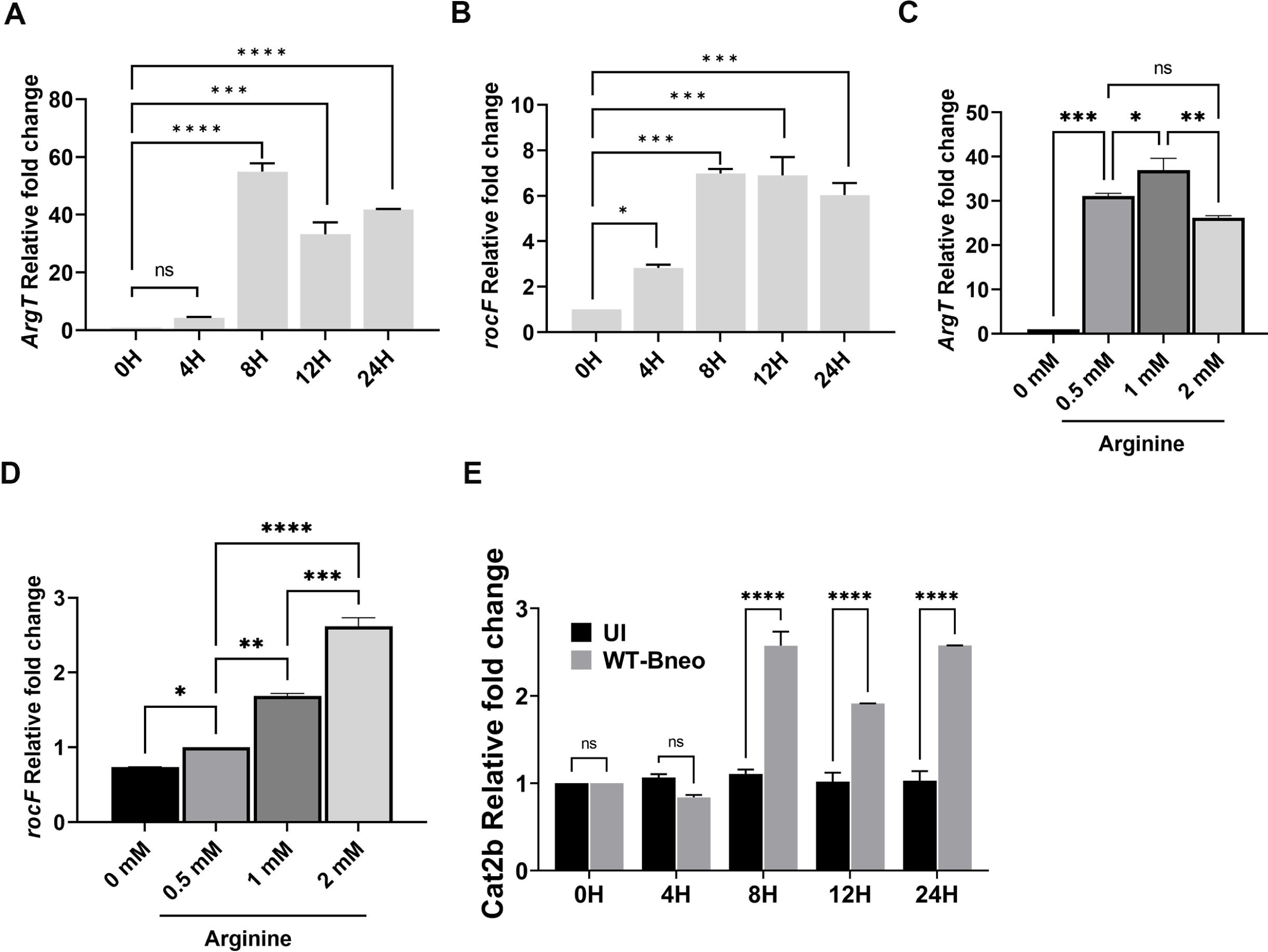
Expression levels of *ArgT* and *rocF* in *B. neotomae* enhanced during infection. **(A, B)** Expression levels of *ArgT* and *rocF* in *B. neotomae* at various time points of infection. RAW264.7 cells were infected with WT-*B. neotomae*, followed by harvesting the infected cells at indicated time points and quantification of expression levels of *ArgT* **(A)** and *rocF* **(B)**of *B. neotomae* by qPCR. **(C, D)** Expression levels of *ArgT* and *rocF* in *B. neotomae* with increasing concentrations of arginine. RAW264.7 cells were infected with WT-*B. neotomae* in DMEM (-) arginine or DMEM (-) arginine supplemented with indicated concentrations of L-arginine for 24 hours, followed by quantification of *ArgT* **(C)**and *rocF* **(D)**expression levels by qPCR. The qPCR data were normalized with 16s rRNA, and relative mRNA expression was quantified with respect to uninfected macrophages for the corresponding time point of infection. **(E)** Upregulation of cationic amino acid transporter in the *Brucella*-infected macrophages. RAW264.7 macrophages were infected with WT-*B. neotomae* (WT-Bneo), followed by quantification of the expression levels of *CAT2b* at indicated time points by qPCR. The data were normalized with GAPDH, and relative mRNA expression was determined with respect to the uninfected macrophages for each time point of infection. The data are presented as the mean ± SEM from at least three independent experiments (*, P <L0.05; **, PL<L0.01; ***, PL<L0.001; ****, PL<L0.0001).

Next, we analyzed whether *Brucella* infection modulates the expression of the host arginine transporter *CAT2b*, which is not constitutively expressed like *CAT1* (*19*). RAW264.7 cells were infected with *B. neotomae*, followed by analysis of the endogenous levels of *CAT2b* by qPCR at various times post-infection. We observed upregulation of *CAT2b* in macrophages at a later time point of infection (Fig. 3E). The experimental data indicated that *CAT2b* is induced in *Brucella*-infected macrophages to promote arginine transport into the infected cells for NO production, as reported for *Salmonella* (16) and *H. pylori* (20) infection.

### The Δ*ArgT*-*B. melitensis* presents an attenuated phenotype in the infected mice

To understand the role of *ArgT in vivo*, we examined the infection dynamics of Δ*ArgT*-*B. melitensis* in mice. Six-to eight-week-old female BALB/c mice were infected with WT or Δ*ArgT B. melitensis*, followed by analysis of the splenic load of *B. melitensis* by enumeration of CFUs at 14-days post-infection. We observed a significant reduction in the splenic load of Δ*ArgT*-*B. melitensis* compared to WT, indicating that Δ*ArgT-B. melitensis* exhibited defective survival in the infected mice (Fig. 4A). To further confirm the data, we examined the spleen and liver weights of mice infected with WT or Δ*ArgT-B. melitensis*. We observed increased spleen and liver weights, and hepatosplenomegaly in mice infected with WT-*B. melitensis* (Fig. 4B-D). In contrast, the spleen and liver appeared normal in the mice infected with Δ*ArgT*-*B. melitensis* (Fig. 4D). Furthermore, we analyzed iNOS expression in the spleen and NO levels in the serum of mice infected with WT or Δ*ArgT*-*B. melitensis*. We found enhanced iNOS expression and potentiated levels of serum NO in mice infected with Δ*ArgT*-*B. melitensis* compared to the WT (Fig. 4E & F). Collectively, our experimental data suggest that Δ*ArgT*-*B. melitensis* was defective in suppressing NO production, leading to enhanced clearance of Δ*ArgT*-*B. melitensis* in infected mice.

**Figure 4.**
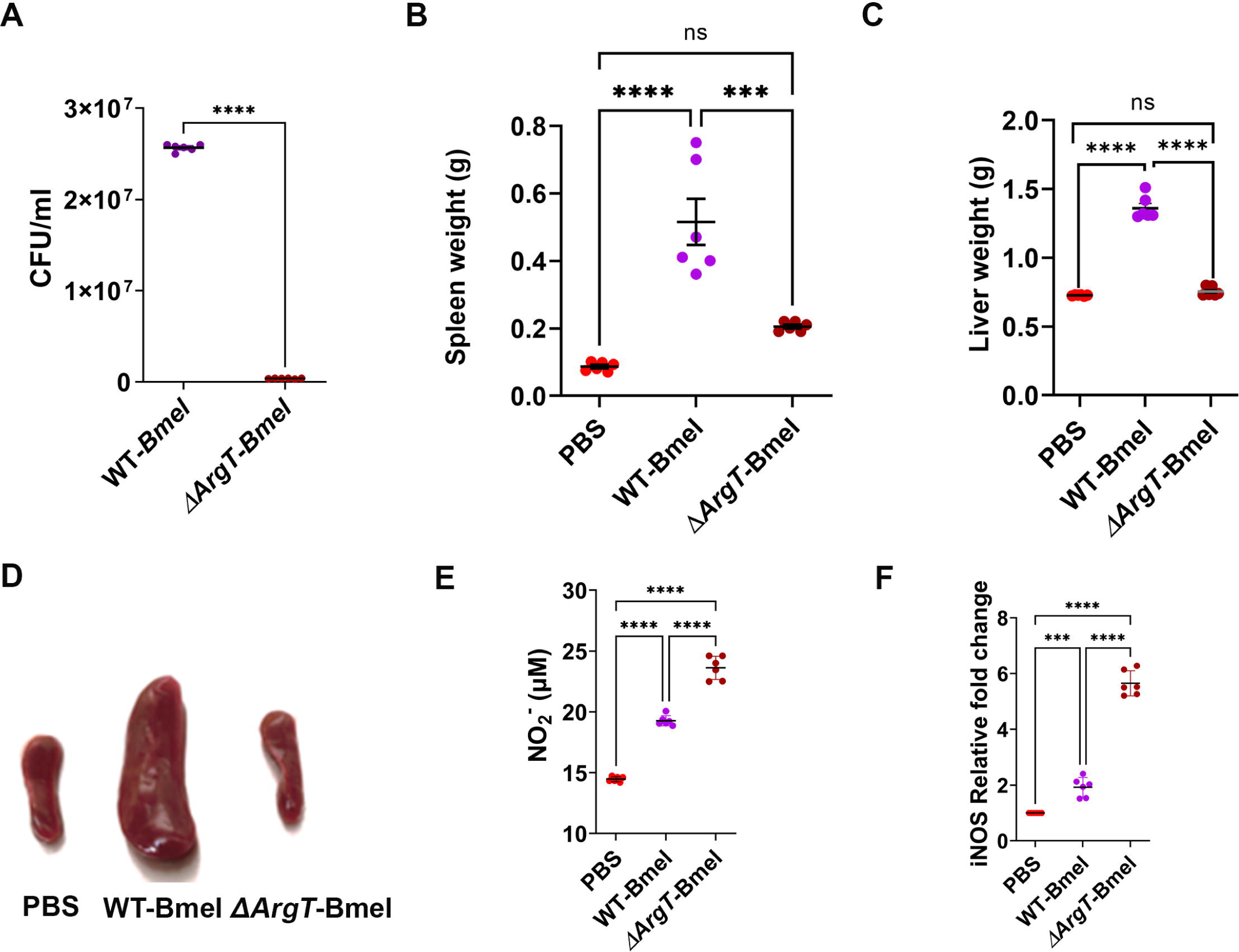
Δ*ArgT-B. melitensis* presents an attenuated phenotype in mice. **(A)** Six to eight weeks old BALB/c mice were infected with WT (WT-BmeI) or Δ*ArgT*-*B. melitensis* (Δ*ArgT*-BmeI). Fourteen days post-infection, the mice were sacrificed, followed by the collection of blood, spleens, and liver. The spleens were homogenized, followed by enumeration of CFU to determine the splenic load of WT or Δ*ArgT*-BmeI. **(A)** The weights of the spleen **(B)** and liver **(C)** of mice infected with WT or Δ*ArgT*-BmeI. **(D)** The photograph showing the spleen of mice infected with WT or Δ*ArgT*-BmeI. PBS was used as the negative control. **(E)** Levels of NO and iNOS**(F)** in the mice infected with WT or Δ*ArgT*-BmeI. The NO levels were determined in the serum of infected mice using Griess reagent, and iNOS expression levels were examined in the spleen by qPCR analysis. The data were normalized with GAPDH, and relative mRNA expression was determined with respect to the group treated with PBS. The data are presented as the mean ± SEM from at least two independent experiments (PL<L0.01; ***, PL<L0.001; ****, PL<L0.0001).

## DISCUSSION

*Brucella* species invade and replicate in various mammalian cell types, including macrophages, dendritic cells, placental trophoblasts, microglia, and epithelial cells (21). The ability of *Brucella* to resist intracellular killing and create a replication-permissive niche contributes to its chronic persistence in an infected host (22). Despite extensive research, minimal information is available on virulence genes and the mechanism by which *Brucella* manipulates cellular processes to survive in the host. Our transposon-based screening identified the arginine/ornithine-binding periplasmic protein precursor *ArgT* (BMEI1022 in *B. melitensis*) as a virulence determinant of *Brucella* and the deletion of *ArgT* affected the survival of *B. melitensis* in macrophages and mice. *ArgT* is reported to be part of the ABC transporter complex involved in importing polar amino acids, where ArgT has been predicted to bind arginine/ornithine (17). Multiple arginine/ornithine-binding periplasmic protein precursors and polar amino acid importers have been annotated in *Brucella* genomes, some of which are absent in particular *Brucella* species. The gene we identified, BMEI1022, was present in all *Brucella* species, except for *B. canis* (17). Furthermore, the arginine/ornithine transporter is present across the microbial kingdom for arginine uptake, which is mainly used as an energy source. Additionally, intracellular bacterial pathogens take advantage of this transporter to deplete arginine from the host cell to neutralize the microbicidal activity of infected cells.

Arginine is a semi-essential amino acid required for protein synthesis that plays a vital role in innate and adaptive immune responses against infectious pathogens (14). Arginine metabolism determines the polarization of macrophages into pro-inflammatory M1 macrophages and anti-inflammatory M2 macrophages (14). In M1 macrophages, arginine is metabolized by NOS to NO and citrulline. There are three isoforms of NOS: inducible NOS is mainly involved in NO production, and its expression is upregulated by various stimulants such as lipopolysaccharide, IFN-γ, TNF-α, and IL-1 (23). The NO free radicals produced by M1 macrophages effectively kill phagocytosed microbial pathogens via nitrosative stress (2). Furthermore, NO combines with superoxide radicals to form other reactive nitrogen species such as peroxynitrite and nitronium ions, contributing to microbial killing (24). M1-polarized macrophages secrete various pro-inflammatory cytokines that play vital roles in the induction of Th1-type adaptive immune responses (14). Another metabolite of arginine, citrulline, can be recycled to arginine for further NO production using the citrulline-NO cycle (25). In contrast, arginase in M2 macrophages converts arginine to ornithine and urea (14). Ornithine is utilized to generate polyamines that regulate various tissue-repairing cellular processes such as DNA replication, cell growth, and differentiation (26). Polyamines have been reported to inhibit the production of proinflammatory cytokines, arginine transport, and NO generation (27). Therefore, M2 macrophages function as anti-inflammatory cells that utilize arginine for tissue repair.

Arginine availability is a rate-limiting step for NO production, and many microbial pathogens exploit this scenario to resist intracellular killing mediated by nitrogen free radicals. One of the microbial strategy to minimize NO production involves the depletion of arginine from infected cells by importing it into microbial cells through the arginine/ornithine transporter. *ArgT* has been reported to play a vital role in the intracellular survival of *S. Typhimurium* where deletion of *ArgT* resulted in enhanced bacterial clearance due to elevated levels of NO (16). Similarly, we found that the deletion of *ArgT* affected the survival of *Brucella* in macrophages and mice. Δ*ArgT*-*Brucella* was defective in importing arginine, as demonstrated by a ^14^C(U)-labeled L-arginine uptake assay. Δ*ArgT*-*Brucella* generates elevated NO levels in infected macrophages and mice, resulting in enhanced clearance.

Since NO production in macrophages depends on the availability of intracellular arginine, it is transported into cells through cationic amino acid transporters (CAT), where CAT1 and CAT2b are highly expressed. *CAT1* is constitutively expressed, whereas *CAT2b* expression is mainly induced by the cytokines IFN-γ, IL-4, or IL-10 (28). The upregulation of *CAT2b* is associated with the induction of NOS and increased NO production in macrophages (29). Enhanced expression of *CAT1* and *CAT2b* and increased uptake of arginine have been reported in *Salmonella*-infected BMDMs and dendritic cells (16). Similarly, we observed upregulation of *CAT2b* in *Brucella*-infected macrophages; however, NO production was minimal in macrophages infected with wild-type *Brucella* compared to those infected with Δ*ArgT*-*Brucella*. This suggests that cytoplasmic arginine is imported by *Brucella* via *ArgT*, which facilitates the depletion of cellular arginine and diminishes NO production.

Arginine metabolism plays a vital role in the microbial pathogenesis. Arginase metabolizes arginine into ornithine and urea, where ornithine is used to synthesize polyamines, which play a vital role in microbial virulence. The microbe-encoded arginase is essential for the virulence of various pathogens including *Leishmania*, *S. Typhimurium*, *H. pylori,* and *M. tuberculosis* (16, 30). We found upregulation of arginase in *Brucella* in culture and infected macrophages. This suggests that *Brucella* can utilize imported arginine to generate polyamines and other metabolites. Polyamines such as spermidine and spermine can suppress the expression of pro-inflammatory cytokines and prevent iNOS induction in infected macrophages (31, 32). Spermidine and spermine induce the expression of virulence genes in bacterial pathogens to facilitate their invasion and intracellular survival (33, 34). Deletion of the arginine transporter results in reduced expression of virulence genes encoding T3SS in Enterohemorrhagic *Escherichia coli* (EHEC), and increased colonic arginine enhances the virulence of *Citrobacter rodentium* (35). The activation of arginase pathways in *Edwardsiella piscicida* inhibits the NLRP3 inflammasome by blocking the efflux of K+ from the macrophages (36). Furthermore, many bacterial pathogens, including *B. abortus*, encode ornithine cyclodeaminase, which converts ornithine to proline and can serve as an essential source of carbon and nitrogen (37). Similarly, host arginase I or arginase II induction has been reported in other microbial pathogens, including *Leishmania major*, *Toxoplasma gondii*, and M. tuberculosis, which enables them to minimize NO production (38, 39). These studies indicate multiple roles for arginine in determining virulence and host responses to infection and serve as a source of nutrition during microbial infection.

The second metabolite, urea, is cleaved into carbon dioxide and ammonia by the microbial urease enzyme, and ammonia can serve as a nitrogen source (40). Furthermore, ammonia generated through the action of arginase can neutralize the acidic pH in phagolysosomes, which can contribute to preventing the intracellular killing of microbial pathogens. *H. pylori* has been reported to generate ammonia, which increases the pH of its surroundings and favors bacterial colonization in the stomach (41). *Brucella* also harbors urease, which can generate ammonia to suppress the microbicidal properties of the infected cells. Therefore, the import of arginine from the host cell through arginine transporters offers multiple benefits to the pathogens, such as the use of arginine as the source of energy, its depletion from the intracellular milieu to limit the production of NO free radicals, and the production of ammonia to neutralize the acidic pH in its microenvironment.

In conclusion, our experimental data indicate that *ArgT* plays an essential role in *Brucella* virulence of *Brucella* in macrophages and mice. It appears that the depletion of intracellular arginine and the conversion of imported arginine into other metabolites, such as polyamines and urea, contribute to the intracellular survival of *Brucella* in infected cells. Furthermore, negative regulation of the M1 polarization of macrophages by minimizing NO production and the resulting suppression of pro-inflammatory cytokines can contribute to the immune evasion of *Brucella*. Our findings provide important insights into the strategies employed by *Brucella* for chronic persistence in infected hosts. Since Δ*ArgT-B. melitensis* presented an attenuated phenotype in macrophages and mice; thus, it can serve as an ideal candidate for developing improved live-attenuated vaccines for brucellosis. Furthermore, *ArgT* could be used to develop novel therapeutic strategies for animal and human brucellosis. These studies also highlight the vital role of arginine in clearing invading microbial pathogens and the strategy of infectious pathogens to successfully overcome this host defense for their survival.

## MATERIALS AND METHODS

### Ethical statement

Six-to eight-week-old female BALB/c mice were obtained from the Animal Resource and Experimental Facility of NIAB, Hyderabad, India. The original breeding colonies were obtained from Jackson Labs (USA). All experiments involving *B. melitensis* were conducted at the Biosafety Level-3/Animal BSL3 facility of UoH-NIAB on the campus of the University of Hyderabad. The experimental protocols were approved by the Institutional Biosafety Committee (Approval number: IBSC/SEPT2019/NIAB/GR01), Institutional Animal Ethics Committee (Approval number: IAEC/NIAB/2022/09/GKR), and BSL-3 Research Review Committee (Approval number: BSL3-Jan2022/003). ARRIVE guidelines have been followed for *in vivo* experiments to examine the attenuation of Δ*ArgT-B. melitensis* in the mouse model of brucellosis.

### Culture of cells and *Brucella*

RAW264.7 murine macrophages (ATCC) were cultured in Dulbecco’s Modified Eagle’s medium (Sigma) supplemented with 10% fetal bovine serum (GIBCO) and 1X penicillin-streptomycin solution (Invitrogen) and maintained at 37 °C in a humidified incubator with 5% CO2. The *B. neotomae* 23459-5K33 was procured from the ATCC. *B. melitensis* 16M was obtained from the ICAR Indian Veterinary Research Institute (Izatnagar, UP, India). *B. neotomae* and *B. melitensis* were cultured on *Brucella* agar or broth (BD Biosciences).

### Generation of transposon insertion mutants of *B. neotomae*

A plasmid harboring the transposable element with a kanamycin resistance cassette, pSAM-Ec, was procured from (Addgene 102939). The pSAM-Ec was introduced into *B. neotomae* by electroporation, and positive colonies were selected on *Brucella* agar plates with kanamycin (40 μg/ml). For the preparation of electrocompetent cells, *B. neotomae* culture with an OD of 0.8-1.0 at 600 nm was collected and kept on ice for 15 min. The culture was then pelleted at 8,000 rpm for 5 min, followed by three washes with ice-cold Milli-Q water. The culture was then resuspended in 100 µL ice-cold water. To perform electroporation, 500 ng of the pSAM-Ec plasmid was added to *B. neotomae* competent cells and incubated on ice for 30 min. The mixture was then transferred to a 1 mm cuvette (ice chilled) for electroporation at 1.5 kV for 5 ms using an electroporator (Bio-Rad). Electroporated samples were mixed with 1 ml of SOC medium and transferred into a culture tube containing 1 ml of *Brucella* broth (BD). Subsequently, cultures were incubated with shaking for 6 h. The culture was then pelleted and resuspended in 100 µL SOC broth, followed by spreading on a *Brucella* agar plate containing kanamycin. The agar plates were incubated for 2-3 days at 37°C with 5% CO2. The colonies obtained were cultured in *Brucella* broth with kanamycin, followed by the preparation of glycerol tubes of cultures, and stored at -80 °C for future use.

### Infection studies using macrophages

RAW264.7 cells were seeded (1х10^5^ cells/well) into 12-well plates without antibiotics and allowed to adhere overnight. Next, the cells were infected with *B. neotomae* at a multiplicity of infection (MOI) of 1000:1 or *B. melitensis* at an MOI of 200:1. After adding *Brucella*, the plates were centrifuged at 280L×Lg for 3 min to pellet the bacteria on the cells. Ninety minutes post-infection, the cells were washed with 1X PBS and treated with gentamycin (30 µg/ml) for 30 min to kill the extracellular bacteria. The cells were then washed with 1X PBS and maintained in 3 µg/ml gentamycin-containing media until harvesting at various times post-infection for CFU analysis. To determine the CFU, the cells were lysed with 0.1% Triton X100 in PBS, followed by plating serial dilutions of the cell lysates onto *Brucella* agar plates with kanamycin. The plates were incubated at 37 °C in the presence of CO2 for 2-3 days. Colonies obtained on the plates were counted and expressed as CFU/ml.

### Identification of transposon insertion site

To identify the transposon insertion site in the genomic DNA of *B. neotomae* mutants that exhibited defective intracellular survival, the genomic DNA of the mutants was isolated using a Wizard Genomic DNA Purification Kit (Promega). The isolated genomic DNA was analyzed by agarose electrophoresis and quantified using a NanoDrop spectrophotometer. Subsequently, 3 µg of genomic DNA was partially digested with *Hha* I enzyme for 10 min. The digested products were purified using the QIAquick PCR Purification Kit (Qiagen) and ligated using T4 DNA ligase (NEB) to obtain circular DNAs containing the kanamycin cassette and the flanking genomic DNA sequences where the transposon insertion has occurred. The ligated products were used as templates for inverse PCR using primers specific for the kanamycin cassette. The PCR products were then cloned into the pCR™4-TOPO® TA cloning vector (Thermo Fisher Scientific) and the clones were confirmed by restriction digestion with *BamH*I. Subsequently, the clones were sequenced using an M13 forward primer to identify the transposon-insertion region.

### Determination of nitric oxide production

Nitrite (NO_2_^−^) released into the culture supernatants of *Brucella*-infected macrophages was used to determine the NO concentration using the Griess Reagent Kit (Invitrogen) according to the manufacturer’s instructions. To measure NO production, macrophages were infected with WT or mutant *B. neotomae* or *B. melitensis,* as described previously. At various times post-infection, the culture supernatants were harvested, mixed with N-[naphthyl] ethylenediamine dihydrochloride (0.01%; 50 µL) and sulfanilamide (0.1%; 50 µL) in 5% phosphoric acid, and incubated for 1 h at RT. The absorbance of the mixture was measured at 540 nm wavelength using a multimode plate reader (PerkinElmer). The concentration of NO was quantified using a standard curve, according to the manufacturer’s instructions.

To analyze NO production upon the addition of arginine, RAW264.7 cells were cultured in arginine-deficient DMEM without phenol red or the same medium supplemented with L-arginine (2 mM), followed by infection with WT or mutant *Brucella* for 24 h. The released nitrite in the culture supernatant was examined using the Griess reagent, as described previously. qPCR was performed using the cDNA prepared from the cells to quantify iNOS expression levels.

### Quantitative RT-PCR analysis of gene expression

Total RNA was isolated from infected macrophages using RNAiso-Plus (Takara), and cDNA was prepared using the Prime Script RT reagent kit (Takara) according to the manufacturer’s protocol. Data were normalized to the endogenous control, GAPDH. To analyze *ArgT* expression, data were normalized to the 16s rRNA of *Brucella*. The primers used for the qPCR analysis are listed in Table 1.

**Table. 1:**
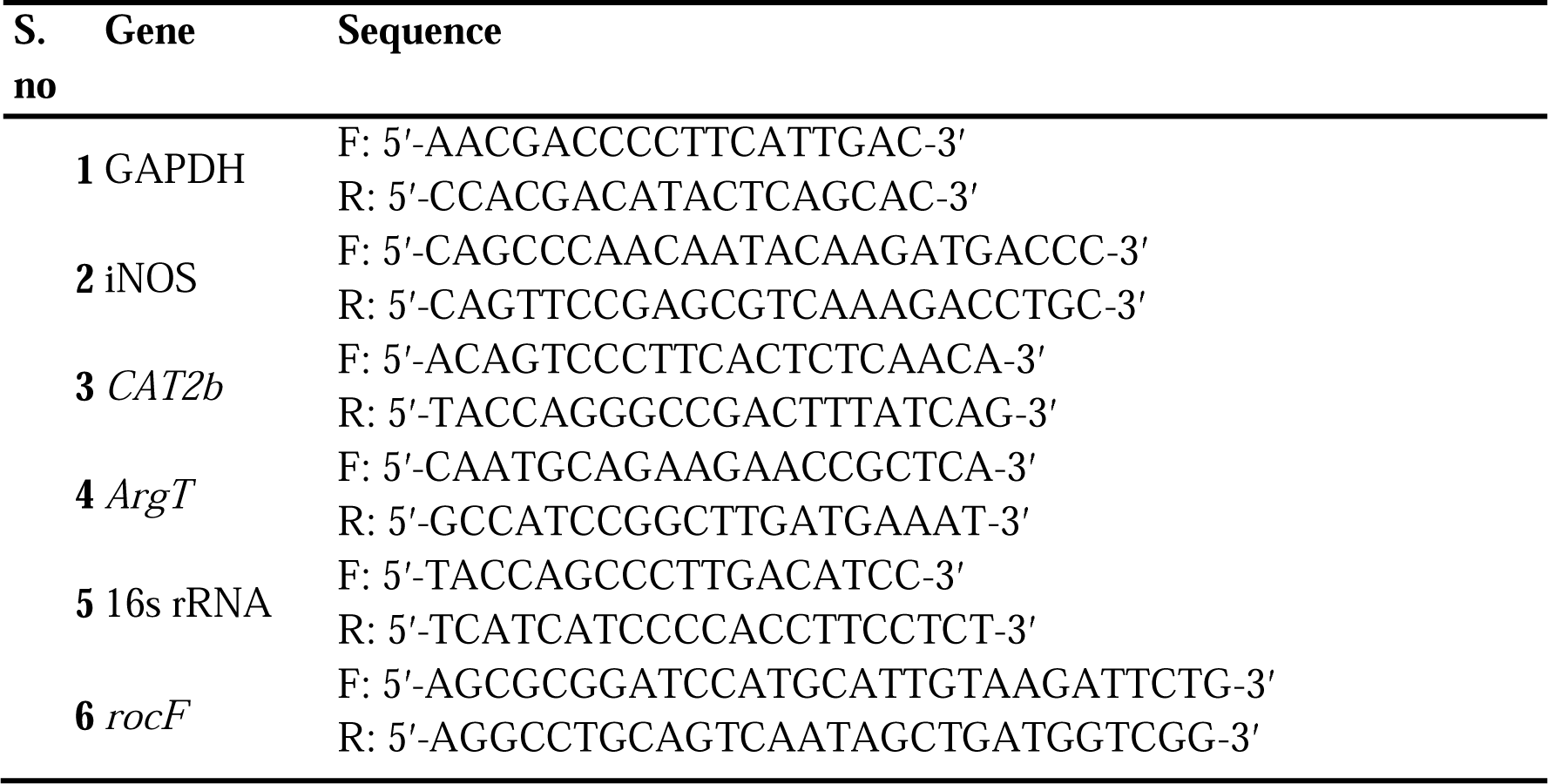
List of primers used for qPCR.

### Generation of ΔArgT-B. neotomae and B. melitensis

Homologous recombination-based gene replacement was used to delete *ArgT* from the chromosomal DNA of *B. neotomae* and *B. melitensis*. To generate the knockout construct, 1kb upstream and 1kb downstream regions of the *ArgT* gene were amplified from the genomic DNA of *B. neotomae* or *B. melitensis*. The forward and reverse primers for amplifying the upstream fragment harbored *Kpn*I and *BamH*I restriction enzyme sites, respectively. Similarly, the forward and reverse primers for amplifying the downstream fragment contained *BamH*I and *Xho*I restriction enzyme sites, respectively. The KanR expression cassette was released from the pUC-kan-LoxP plasmid using *BamH*I restriction enzyme. Subsequently, these three fragments were purified and ligated into *Kpn*I and *Xho*I sites of the pZErO1 vector (Thermo Fisher Scientific) using three-way 3-way ligation. The resulting *ArgT* knock-out construct was confirmed by restriction enzyme digestion and sequencing. Subsequently, the *ArgT* knockout construct was electroporated into *B. neotomae* or *B. melitensis,* as described previously. The upstream and downstream fragments in the KO plasmid recombined with the respective sequences on the chromosomal DNA of *B. neotomae,* replacing the *ArgT* gene with the KanR expression cassette. The transformed *B. neotomae* or *B. melitensis* colonies growing on kanamycin and zeocin, which indicated a single recombination event and the resulting insertion of the KO plasmid into the chromosomal DNA, were discarded. Zeocin-sensitive and kanamycin-resistant Δ*ArgT*-*B. neotomae* or *B. melitensis* colonies were selected and confirmed using PCR.

### Complementation with Δ*ArgT Brucella* with *ArgT* expression plasmid

Complement Δ*ArgT-B. neotomae* or *B. melitensis* with *ArgT* expression plasmid, *ArgT* was cloned into the *Brucella* expression plasmid pNSTrcD at the *BamH*I and *Xho*I sites, respectively. Subsequently, pNSTrcD-*ArgT* was introduced into *B. neotomae* or *B. melitensis* through electroporation, and transformants were selected on *Brucella* agar with chloramphenicol (40 µg/ml).

### Immunoblotting

The cells were lysed in radioimmunoprecipitation assay (RIPA) buffer containing a protease inhibitor cocktail (Pierce). The total protein content of the samples was quantified using Bradford assay (Sigma). Equal concentrations of protein samples were mixed with 5x Laemmle buffer (Bio-Rad), and the samples were boiled for 10 min at 100 °C. Next, the protein samples were run on an SDS-PAGE gel, followed by transfer onto a polyvinylidene difluoride (PVDF) membrane using a wet tank blotting system (Bio-Rad). The membrane was blocked with 5% skimmed milk in Tris-buffered saline with Tween-20 (TBST; Cell Signalling Technology) for 1 h, followed by incubation with the respective primary antibody overnight at 4 °C, washing three times with TBST, and incubation with horseradish peroxidase (HRP)-conjugated secondary antibody. Primary or secondary antibodies were diluted in 5% skim milk in TBST. Finally, the membrane was washed three times with TBST and incubated with SuperSignal West Pico or Femto chemiluminescent substrate (Pierce). The signals were captured using a chemi-documentation system (Syngene). The antibody dilutions used are listed in Table 2.

**Table. 2:** List of antibodies used in the study.

| S. no | Antibody | Source | Dilution |
| --- | --- | --- | --- |
| 1 | Inos | Cell Signalling Technology | 1:1000 |
| 2 | Anti-actin HRP-conjugated | Sigma Aldrich | 1:25,000 |
| 3 | Anti-rabbit HRP-conjugated | Cell Signalling Technology | 1:5,000 |

### Generation of *B. neotomae* expressing GFP for fluorescence microscopy

To generate *B. neotomae* expressing green fluorescent protein (GFP), *Brucella* expression plasmid (pNSTrcD) harboring GFP was electroporated into wild-type or mutant *B. neotomae* as described previously. Transformants were selected on a *Brucella* agar plate containing chloramphenicol (40 µg/mL). For fluorescence microscopy, RAW264.7 cells were seeded into a glass-bottom imaging dish (5 × 10^4^ cells/dish) and allowed to adhere overnight. Next, cells were infected with WT or mutant *B. neotomae*-GFP for 24 h. Subsequently, the infected cells were fixed with 4% paraformaldehyde in PBS, and the nuclei were stained with Hoechst stain (Thermo Fisher Scientific). Images were captured using a fluorescence microscope (Carl Zeiss) at 20X magnification. Twelve fields were examined for each sample.

### Arginine uptake assay

The ability of the wild-type or Δ*ArgT*-*B. neotomae* to uptake arginine was determined using radioactive ^14^C(U)-L-arginine. To analyze this, wild-type or Δ*ArgT* or Δ*ArgT*+*ArgT*-*B. neotomae* were cultured in arginine-deficient DMEM. Subsequently, the cultures were harvested at stationary phase (1.0 OD at 600 nm), followed by pelleting down the bacteria and washing the pellet thrice with pre-warmed uptake solution (137 mM NaCl, 5.4 mM KCl, 1.2 mM of MgSO4.7H2O, 2.8mM of CaCl2.2H2O, 10mM of HEPES and 1mM of KH2PO4, pH.7.4). The bacterial pellets were resuspended in 100 µL of pre-warmed uptake solution containing 10 mM of L-leucin and 100 µM of ^14^C(U) L-arginine (0.1 mCi/ml) and incubated for 1 h at 37 °C in an incubator with 5% CO2. Next, the bacterial pellets were washed three times with the stop solution (10 mM HEPES, 10 mM TRIS, 137 mM NaCl, and 10 mM non-radioactive L-arginine; pH 7.4) to remove the unincorporated radiolabeled ^14^C(U)-L-arginine. Subsequently, the bacterial pellets were lysed in B-PER (Pierce), and the uptake of ^14^C(U) l-arginine was measured in the bacterial lysates using a liquid scintillation counter (Perkin Elmer).

To examine the incorporation of radiolabeled-arginine into *Brucella* proteins, an uptake assay was performed with ^14^C(U) L-arginine, as described above. The bacterial pellets were resuspended in *Brucella* broth and cultured for 24 h. Bacteria were pelleted by centrifugation and lysed using B-PER. The total protein concentration in the lysates was quantified using Bradford reagent (Sigma), and equal amounts of bacterial proteins were resolved by SDS-PAGE. Subsequently, the SDS-PAGE gel was wrapped with silicon, kept in an autoradiography cassette for 24 h, and analyzed using a phosphor-imager (Amersham) to measure the incorporated ^14^C(U)-L-arginine into the total protein of wild-type, Δ*ArgT or* Δ*ArgT*+*ArgT*-*B. neotomae*.

### Mice infection

Six-to seven-week-old female BALB/c mice (6 mice/group; Table 3) were intraperitoneally injected with WT or Δ*ArgT*-*B. melitensis* (2×10^6^ CFU/mouse). Fourteen days post-infection, the mice were euthanized, and their livers and spleens were collected aseptically (Table 4). Spleen and liver weights were measured using a digital weighing balance. For CFU enumeration, spleens were homogenized in 1X PBS (1 ml) using a bead beater, followed by clarifying the homogenates by centrifugation. Subsequently, the homogenates were serially diluted in 1X PBS and plated on *Brucella* agar plates. The plates were incubated at 37° C with 5% CO2 for 2-3 days, followed by enumeration of the CFU. To examine iNOS levels, mice (5 mice/group) were infected with WT or mutant *B. melitensis* as described previously. The spleens were collected four days post-infection, followed by total RNA isolation and cDNA preparation. Subsequently, iNOS levels were determined by qPCR analysis. Serum isolated from the blood of infected or control mice was used to analyze NO production using Griess reagent.

**Table 3.**
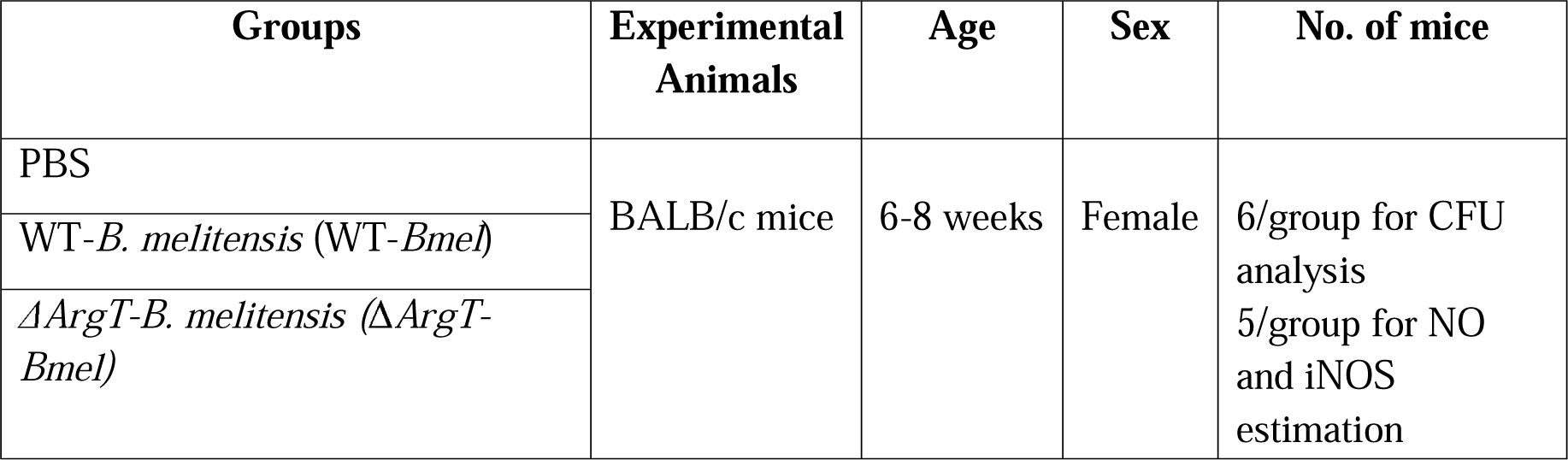
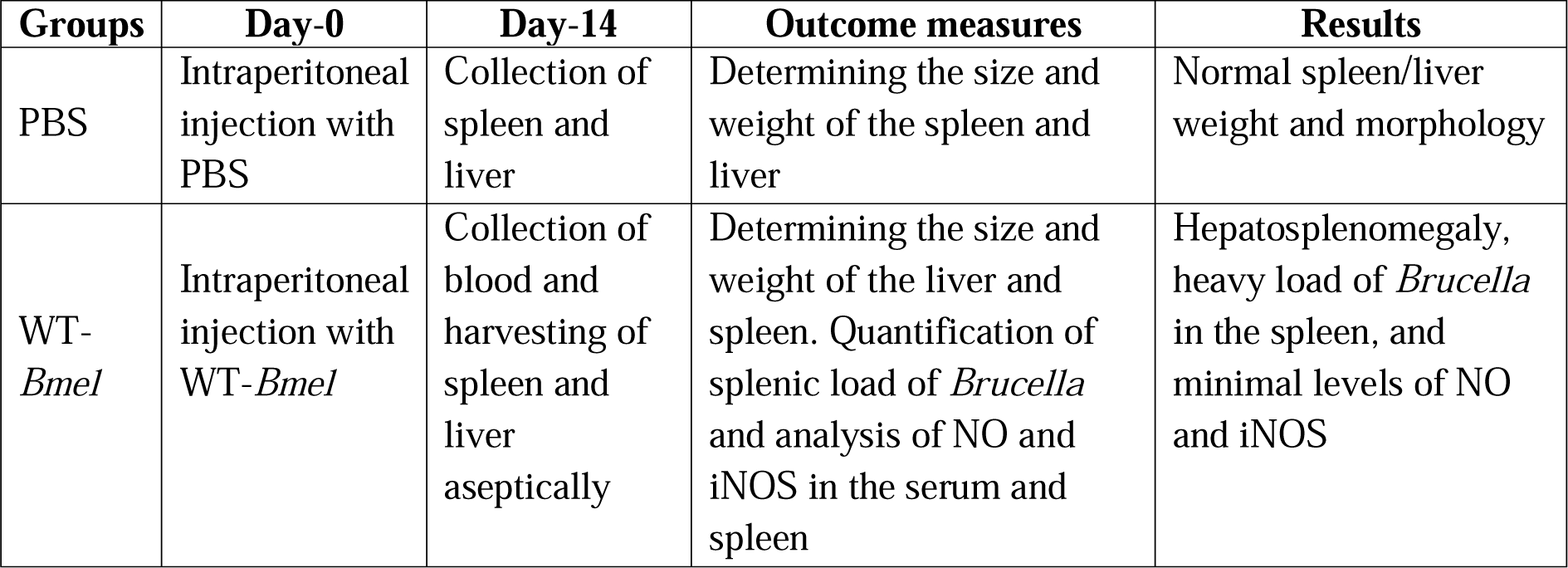
Details of experimental animals to examine the attenuation of Δ*ArgT-B. melitensis*.

**Table 4.** Study design to examine the attenuation of Δ*ArgT-B. melitensis* in the mice model of brucellosis.

|  |  |  |  |  |
| --- | --- | --- | --- | --- |
| $\Delta ArgT$-*Bmel* | Intraperitoneal injection with $\Delta ArgT$-*Bmel* | Collection of blood and harvesting of spleens and liver aseptically | Determining the size and weight of the liver and spleen. Quantification of splenic load of *Brucella* and analysis of NO and iNOS in the serum and spleen | No hepatosplenomegaly, minimal load of *Brucella* in the spleen, and enhanced levels of NO and iNOS |

### Statistical analysis

Data are shown as the mean of triplicate measurements ± standard error of the mean (SEM). The level of significance was set at P < 0.05. GraphPad Prism software (version 7.0) was used for statistical analysis of the experimental data. Data are presented as mean ± standard error of the mean (SEM). Statistical significance was determined by t-tests (two-tailed) for pairwise comparisons. One-way analysis of variance (ANOVA) Dunnett’s hypothesis test was used to analyze data involving more than two samples.

## ACKNOWLEDGMENTS

We thank the National Institute of Animal Biotechnology for providing funding and experimentation. We acknowledge the BSL3/A-BSL3 facility of UoH-NIAB for the experiments with *B. melitensis*. The SRM acknowledges a research fellowship from the University Grant Commission (UGC).

## ABBREVIATION

iNOS: Induced nitric oxide synthase.
NO: Nitric Oxide
Cat: Cationic amino acid transporter
*ArgT*: Arginine ornithine binding transporter protein
Δ*ArgT*: Arginine ornithine binding transporter protein knockout.
rocF: Arginase

## AUTHOR’S CONTRIBUTION

GR conceived and designed the study. SRM and KJ performed the experiments and analyzed the experimental data. GR and SRM prepared the manuscript. All authors have reviewed the results and approved the final version of the manuscript.

